# Secretion, maturation and activity of a quorum-sensing peptide (GSP) inducing bacteriocins transcription in *Streptococcus gallolyticus*

**DOI:** 10.1101/2020.06.04.133652

**Authors:** Anthony Harrington, Alexis Proutiere, Shaynoor Dramsi, Yftah Tal-Gan

## Abstract

*Streptococcus gallolyticus* subsp. *gallolyticus* (*Sgg*) is an emerging opportunistic pathogen responsible for septicemia and endocarditis in the elderly. Invasive infections by *Sgg* are strongly linked to the occurrence of colorectal cancer (CRC). It was previously shown that increased secondary bile salts in CRC-conditions enhances the bactericidal activity of gallocin, a bacteriocin produced by *Sgg*, enabling it to colonize the mouse colon by outcompeting resident enterococci. In a separate study, we have shown that *Sgg* produces and secretes a 21-mer peptide that activates bacteriocin production. This peptide was named CSP because of its sequence similarity with competence stimulating peptides found in other streptococci. Here we demonstrate that CSP is a *bona fide* quorum-sensing peptide involved in activation of gallocin gene transcription. We therefore refer to CSP as GSP (gallocin stimulating peptide). GSP displays some unique features since its *N*-terminal amino-acid lies three residues after the double glycine leader sequence. Herein, we set out to investigate the processing and export pathway that leads to mature GSP. We also conducted the first comprehensive structure-activity relationship (SAR) of *Sgg* GSP to identify its key structural features.

**Significance:** *Streptococcus gallolyticus* subsp. *gallolyticus* (*Sgg*) is an opportunistic pathogen associated with colorectal cancer (CRC) and endocarditis. *Sgg* utilizes quorum-sensing (QS) to regulate the production of a bacteriocin (gallocin) and gain selective advantage in colonizing the colon. In this manuscript, we report 1) the first structure-activty relationship study of the *Sgg* QS pheromone that regulates gallocin production; 2) evidence that the active QS pheromone is processed to its mature form by a unique ABC transporter and not processed by an extracellular protease; and 3) supporting evidence of interspecies interactions between streptococci pheromones. Our results revealed the minimal pheromone scaffold needed for gallocin activation and uncovered unique interactions between two streptococci species QS signals that warrant further studies.

## Introduction

*Streptococcus gallolyticus* subsp. *gallolyticus* (*Sgg*), previously known as *Streptocococcus bovis* biotype I, is an emerging opportunistic human pathogen belonging to the highly diverse *Streptococcus bovis*/*Streptococcus equinus* complex (SBSEC) (1-3). *Sgg* is responsible for causing infective endocarditis, septicemia and has been consistently associated with colorectal cancer (CRC) (4, 5). Recent experimental data support both a passenger and driver role of *Sgg* in CRC development (6-8). Using a murine CRC model, it was shown that CRC-specific conditions strongly promote colonization of the colon by *Sgg* (9). Indeed, increased levels of secondary bile salts in tumor-bearing mice enhanced the bactericidal activity of gallocin, a putative class IIb bacteriocin encoded by two genes *blpA and blpB*, produced by *Sgg*, thus enabling it to colonize the mouse colon by outcompeting resident enterococci.

It was later reported that *Sgg* strain TX20005 secretes a 21-mer peptide that induces the production of unknown bacteriocins that are active against various oral streptococci (10). This peptide was named competence stimulating peptide (CSP) because of its sequence similarity with other CSPs found in other streptococci. However, under the conditions tested, *Sgg* CSP was unable to induce natural competence as measured by plasmid DNA uptake (10).

Originally discovered in *Streptococcus pneumoniae*, natural competence was shown to be a tightly regulated process involving a hormone-like cell product, termed pheromone (11). The nature of the molecule inducing competence in pneumococci was identified as a linear unmodified 17-residue peptide named CSP (for competence stimulating peptide) (12). CSP, encoded by *comC*, is synthesized as a precursor peptide of 41 residues containing the Gly-Gly consensus processing site found in peptide bacteriocins (13). CSP is secreted and maturated by a specialized ATP-binding transporter, ComAB, that cleaves its *N*-terminal part just after the Gly-Gly motif (14). Once CSP has reached a threshold concentration in the extracellular medium, it binds to a transmembrane histidine kinase receptor, ComD, which in turn triggers phosphorylation of ComE, a response regulator that activates the transcription of *comX*, the master regulator of competence genes (14, 15). *comX*/*sigX* encodes an alternative sigma factor allowing the coordinated expression of a set of approximately 20 genes encoding the competence machinery. This *comABCDE* quorum sensing (QS) circuitry has been found to regulate competence in 12 streptococci species belonging to the *Mitis* and *Anginosus* groups (16-18). In 2010, two groups showed that natural competence could be induced in a wide range of streptococci by a second QS circuitry involving a 7-amino acid peptide called XIP (for *sigX*/*comX* inducing peptide) (19, 20).

Importantly, competence and bacteriocin production in streptococci are often coupled processes that are regulated differentially by the two QS circuitries described above (21). For example, the *comABCDE* circuitry in *S. mutans* regulates the production of bacteriocins called mutacins directly and competence through activation of the *comRS* circuitry (14, 22-25). This observation has led to the proposal that the *S. mutans* CSP be renamed to mutacin inducing peptide (MIP) (14, 26, 27). Additionally, it was shown that the secreted 21-mer CSP/MIP peptide is inactive and requires further processing by the streptococcal extracellular protease (SepM), which cleaves the three *C*-terminal residues of 21-mer CSP/MIP to generate the active 18-mer CSP/MIP (28, 29).

*Sgg* possess both the *comABCDE* and *comRS* loci in its genome (30-32). In the accompanying paper, it is demonstrated that *Sgg* CSP 24-mer is a *bona fide* QS inducing peptide involved in activation of gallocin transcription (Proutière et al., 2020). However, despite several attempts, we failed to induce competence in *Sgg* with CSP (Harrington et al. 2018, Proutière *et al*., 2020) (9, 10). We therefore propose referring to *Sgg* CSP as GSP (gallocin stimulating peptide) and to renamed its putative associated genes *comAB* as *blpT1-2* (bacteriocin like peptide transporter) and *comDE* as *blpHR* (bacteriocin like peptide histidine kinase and regulator, respectively). Since GSP is predicted to be a 24-mer peptide while the isolated GSP was found to be a 21-mer peptide starting three residues after the double glycine leader sequence of the precursor peptide, we first tested the activity of GSP 24-mer versus GSP 21-mer on gallocin transcription using a green fluorescent protein (GFP) reporter construct described in the accompanying paper (Proutière *et al*., 2020). Next, we set out to explore how GSP was processed and evaluated the potential role of *Sgg* extracellular protease, which is homologous to SepM. Finally, we undertook a comprehensive structure function analysis of the GSP pheromone, which revealed that almost half of the peptide sequence (first nine *N*-terminal residues) is dispensable and can be removed without significantly affecting the GSP ability to activate its cognate histidine kinase receptor, providing a minimal structure for the development of GSP-based QS inhibitors that could affect *Sgg* fitness during competition with the gut microflora.

## Results & Discussion

### GSP active form is a 21-mer peptide

We first aimed at determining the active form of GSP in *Sgg* strain UCN34. GSP is derived from a 45 residue precursor encoded by *gsp* gene, and is predicted to be processed to a 24-mer peptide by cleavage of the amino terminal leader sequence after a double glycine motif, then exported to the extracellular environment by the BlpT transporter. However, GSP was previously isolated as a 21-mer peptide from *Sgg* TX20005 (10). We first confirmed by mass spectrometry analysis that GSP isolated from culture supernatants of strain UCN34 was strictly identical to the 21-mer peptide previously purified from strain TX20005 (**Fig. S1)**. The predicted 24-mer GSP was shown to activate transcription of the gallocin genes (Proutière et al., accompanying paper). Therefore, we chemically synthesized GSP 21-mer and GSP 24-mer peptides to test their efficiency in activating gallocin gene transcription. To do so, we used a reporter *Sgg* UCN34 strain in which the endogenous *gsp* gene has been deleted and containing a plasmid expressing *gfp* under the control of the gallocin genes promoter (P*blpAB*) (Proutière et al., accompanying paper). No expression of *gfp* was observed in the absence of GSP whereas both 24-mer and 21-mer GSP peptides were able to activate *blpAB* gene transcription. The GSP 21-mer was found to be more active than the 24-mer, as can be seen from the EC_50_ values of both peptides (i.e. the concentration of peptide needed to reach 50% of maximal receptor response). Indeed, GSP 21-mer peptide EC_50_ (2.96 nm) was around 100 times lower than GSP 24-mer EC_50_ (287 nM), indicating that the 21-mer is significantly more active than the 24-mer (**Table 1**). These results imply that GSP maturation to a 21-mer peptide increases GSP efficiency as compared to the 24-mer peptide, but is not absolutely required, unlike the CSP/MIP of *S. mutans* (28, 29).

**Table 1.** EC_50_ values of *Sgg* GSP 21-mer and 24-mer in UCN34Δ*gsp*^a^

| Compound | Sequence | EC<sub>50</sub> (nM)<sup>b</sup> | 95% CI<sup>c</sup> |
| --- | --- | --- | --- |
| GSP 21-mer | DFLIVGPFDWLKKNHKPTKHA | 2.96 | 1.70 – 5.14 |
| GSP 24-mer | KNKDFLIVGPFDWLKKNHKPTKHA | 287 | 145 – 568 |

<sup>a</sup> See Materials and Methods for details of reporter strains and methods. All assays were performed in triplicate. <sup>b</sup> EC<sub>50</sub> values were determined by testing peptides over a range of concentrations. <sup>c</sup> 95% confidence interval.

### An ABC transporter is responsible for the secretion of GSP and gallocin peptides

We next aimed at determining how GSP was secreted and processed into a 21-mer peptide in *Sgg* UCN34. In *S. mutans*, the equivalent 21-mer MIP peptide encoded by *comC* is secreted by a specific ABC transporter composed of ComA and ComB proteins (**Fig. 1A**). Two genes (*gallo_rs10390/10395*) encoding a putative ABC transporter homologous to ComA/B were found in the vicinity of the GSP and gallocin genes (**Fig. 1A**). To test the role of this putative transporter in GSP secretion and maturation, an in-frame deletion mutant of *gallo_rs10390/10395* was constructed in strain UCN34 and will be referred to as UCN34:Δ*blpT*. For each deletion mutant generated in *Sgg* clinical isolate UCN34, we also selected a clone that reverted to the WT genotype (bWT) following homologous recombination. These bWT strains should display the WT phenotype and are isogenic to their mutant counterparts. Basic phenotypic characterization of several clones for a given mutation was carried out systematically (growth curve, immunofluorescence microscopy, antibiotic resistance profile) to rule out any major secondary mutations that may have occurred during the engineering process. We then determined gallocin activity against a highly sensitive bacteria, *Streptococcus gallolyticus* subspecies *macedonicus* (*SGM*) using the supernatants of the Δ*blpT* mutant compared to its bWT strain. As shown in **Fig. 1B**, gallocin activity was completely absent in the Δ*blpT* mutant supernatant while present in the bWT. Three possibilities can explain these results: i) absence of GSP in the supernatant of Δ*blpT* mutant; ii) absence of the two structural peptides (BlpA/B) constituting the active gallocin in the supernatant of Δ*blpT* mutant; or iii) absence of both GSP and gallocin peptides in the supernatant of Δ*blpT* mutant. To differentiate between these alternatives, we first tested the ability of the Δ*blpT* supernatant to activate *gfp* transcription in the UCN34 Δ*gsp* pTCVΩP*blpAB*-*gfp* reporter strain mentioned above. To do so, we resuspended the reporter strain in different supernatants. If GSP is present in the supernatant, the gallocin promoter will be activated, turning on GFP and the bacteria will be fluorescent. As shown in **Fig. 1C**, the reporter strain is not fluorescent in the absence of GSP (THY only and Δ*gsp* supernatant) and becomes fluorescent in the presence of GSP (THY + GSP and WT supernatant). No GFP fluorescence could be detected using the Δ*blpT* supernatant, while a strong signal was detected in the bWT supernatant, demonstrating that GSP is not secreted in the Δ*blpT* mutant, an observation that was verified by LC-MS (**Fig. 1C** and **Fig. S2**). These results were further confirmed by introducing the reporter plasmid (pTCVΩP*blpAB*-*gfp*) into the Δ*blpT* mutant and measuring GFP fluorescence. Gallocin promoter was inactive in the Δ*blpT* pTCVΩP*blpAB*-*gfp* strain, while addition of synthetic GSP in the culture medium restored full gallocin promoter activity (**Fig. 1D**).

**Figure 1.**
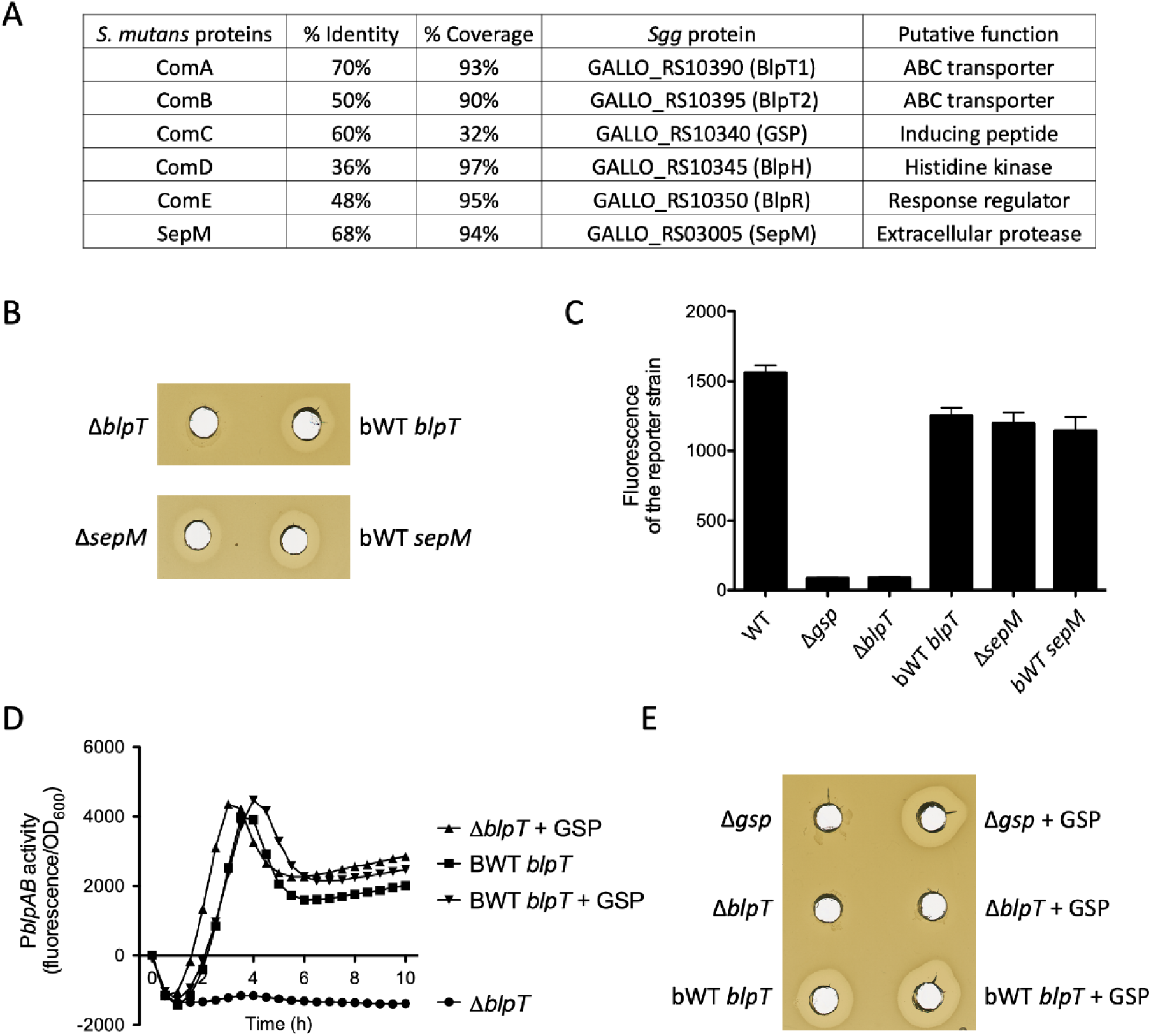
The ABC transporter BlpT secretes both GSP and gallocin peptides. **A**: Summary table of blast results comparing the ComABCDE system and SepM proteins of *S. mutans* with their homologous counterpart in *Sgg*. **B**: Agar diffusion assay showing gallocin activity in the culture supernatant against *Sgm*. Strain tested: *Sgg* UCN34 Δ*blpT* and Δ*sepM* and their bWT counterpart. **C**: Fluorescence of the reporter strain *Sgg* UCN34 Δ*gsp* pTCVΩP*blpAB*-*gfp* resuspended in the supernatant of *Sgg* UCN34 WT, Δ*gsp*, Δ*blpT*, bWT *blpT*, ΔsepM and bWT *sepM*. The fluorescence of the reporter strain increases only if the supernatant contains GSP. **D**: P*blpAB* activity in *Sgg* UCN34 Δ*blpT* and its bWT counterpart with or without addition of 20 nM synthetic GSP. **E**: Agar diffusion assay showing gallocin activity in the culture supernatant against *Sgm*. Strain tested: *Sgg* UCN34 Δ*gsp*, Δ*blpT* and bWT *blpT* cultivated with or without addition of 20 nM synthetic GSP.

We next investigated whether the same ABC transporter (named BlpT) could also secrete the gallocin BlpA/B peptides, and thus analyzed gallocin activity in the Δ*blpT* mutant in the presence of synthetic GSP. As shown in **Fig. 1D**, addition of GSP in the culture medium restored gallocin production in the control Δ*gsp* mutant. However, no gallocin activity could be detected in the Δ*blpT* mutant even in the presence of synthetic GSP, although the gallocin promoter was fully active under these conditions as mentioned above. Altogether, these results indicate that the ABC transporter secretes both GSP and the gallocin peptides.

### SepM is not involved in GSP maturation in *Sgg* UCN34

In *S. mutans* an extracellular protease, SepM, is responsible for CSP/MIP maturation by cleaving the three *C*-terminal amino acids of ComC after its secretion. Blast analysis revealed that *Sgg* UCN34 possess a close SepM homolog (**Fig. 1A**). Therefore, we reasoned that *Sgg* SepM could be involved in GSP maturation by cleaving the three *N*-terminal residues of GSP 24-mer after secretion by the ABC transporter. To test this hypothesis, we deleted *sepM* in *Sgg* UCN34. As shown in **Fig. 1B**, gallocin production appears slightly reduced in the Δ*sepM* mutant as compared to the isogenic bWT. To test whether the Δ*sepM* mutant was able to produce the fully active GSP 21-mer, we first analyzed the capacity of Δ*sepM* supernatants to induce gallocin promoter activity in the reporter strain Δ*gsp* pTCVΩP*blpAB*-*gfp*. We found that the Δ*sepM* supernatant was able to fully activate P*blpAB* promoter in the reporter strain, suggesting that Δ*sepM* supernatants possess the active form of GSP. We then further assessed the Δ*sepM* supernatant using LC-MS and detected the presence of the mature 21-mer GSP (**Fig. S3**). Together, our results indicate that SepM is not involved in GSP maturation in UCN34.

To further evaluate if a cell-bound protease is involved in GSP maturation, we incubated KNK-GSP (24-mer) and GSP (21-mer) with washed UCN34 Δ*gsp* or Δ*sepM* cells in sterile saline solution for 30 minutes and analyzed the filtrates using LC-MS. Degradation of KNK-GSP to GSP was not observed following incubation with either UCN34Δ*sepM* or UCN34Δ*gsp* (**Figs. S4-S5**). Both GSP and KNK-GSP were degraded into GSP-des-D1-L3, suggesting that another cell-bound protease exists in *Sgg* (**Figs. S4-S6**). We hypothesize that in *Sgg*, the specific BlpT ABC transporter has a unique feature that allows it to process GSP to the mature GSP 21-mer, which if true will challenge the double glycine cleavage rule.

Biswas and co-workers have previously shown that the SepM proteases are quite promiscuous and can process CSP signals of other species (28). We wanted to confirm that the *Sgg* SepM is a functional protease and repeated the cell-washed processing assays described above, but this time added the *S. mutans* 21-CSP to washed UCN34Δ*gsp* or UCN34Δ*sepM* cells. The 18-CSP was found in filtrates treated with UCN34 Δ*gsp* but not in filtrates treated with UCN34 Δ*sepM* indicating that *Sgg* SepM is functional and capable of processing *S. mutans* inactive 21-CSP into active 18-CSP **(Figs. S7-S8**) (28). Our results therefore support Biswas findings that SepM found in some streptococci can process *S. mutans* 21-mer CSP into 18-mer CSP. Lastly, we tested 21-mer *Sgg* GSP with *S. mutans* Δ*comC* washed cells. Surprisingly, we observed that *S. mutans* is capable of processing GSP into GSP des-D1-L3 (**Fig. S9**). Combined, our results indicate potential interspecies QS interactions between *Sgg* and *S. mutans*, although their exact nature is currently unclear.

### Structure-Activity Relationships of *Sgg* GSP

Our second aim in this study was to evaluate and determine the key structural motifs that drive GSP binding to its cognate histidine kinase, BlpH, thus activating gallocin gene transcription.

### Alanine scan of *Sgg* GSP

We first set out to perform a full alanine scan of the 21-mer GSP signal. Peptides were synthesized using standard solid-phase peptide synthesis (SPPS) conditions using a CEM Liberty1 microwave synthesizer on Cl-MPA Protide resin (LL). The peptides were then purified to homogeneity (>95%) using RP-HPLC and validated using Mass Spectrometry (for full experimental details and peptide characterization, see Supporting Information). The alanine scan revealed several residues that are important for receptor binding, as mutating them to alanine resulted in significant reduction in activity, yet no one residue was found to be critical for receptor activation, leading to the identification of a competitive inhibitor (**Table 2**). Specifically, the alanine scan revealed that the *C*-terminal half of the peptide (residues W10 – H20) was more important for receptor binding compared to the *N*-terminal half (residues D1 – D9), as modifications to the *C*-terminal half resulted in a significant decrease in potency, with the most important residue being W10 (**Table 2**).

**Table 2.**
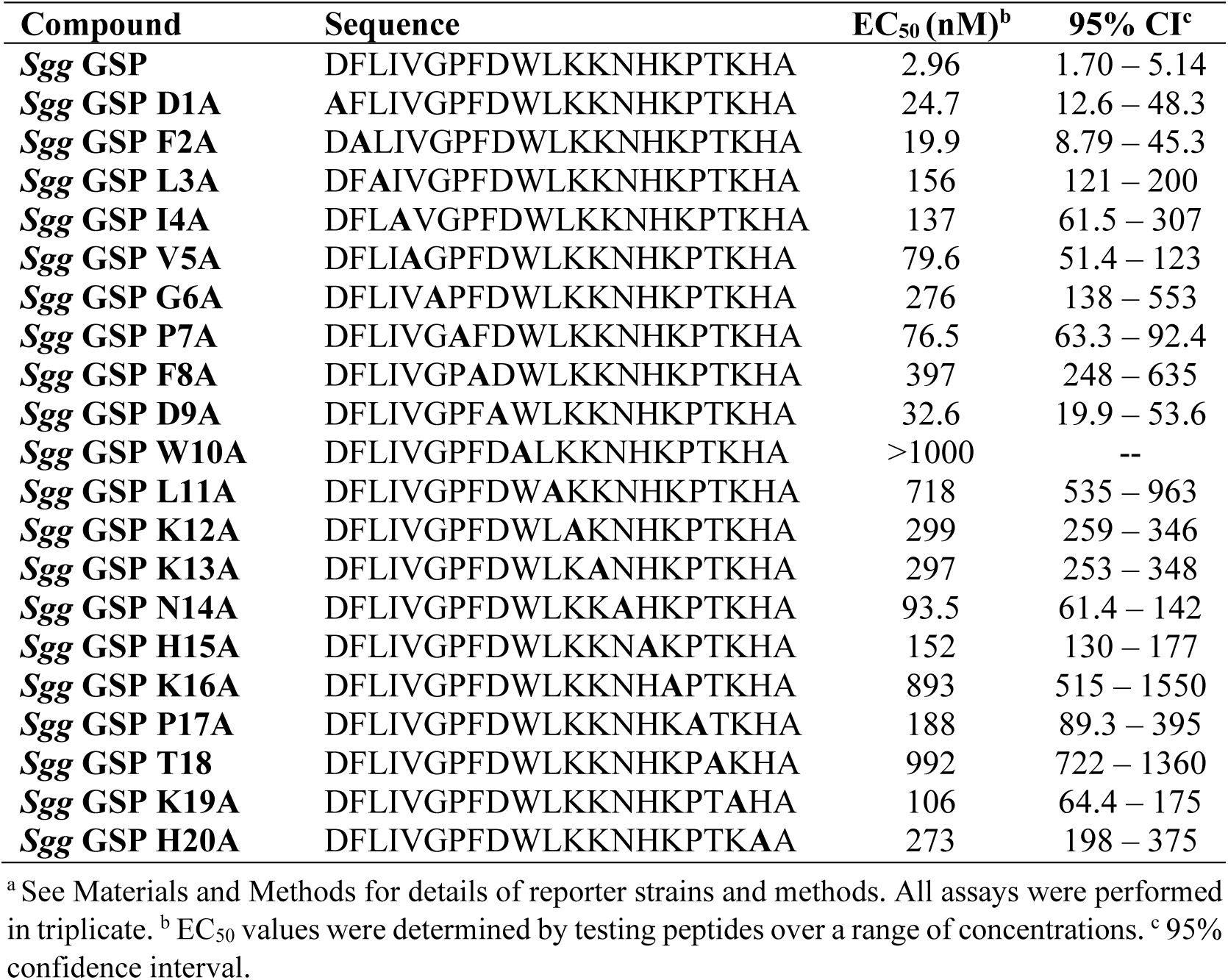
EC_50_ values of *Sgg* GSP alanine analogs in UCN34Δ*gsp*^a^

### Truncation studies of *Sgg* GSP

To further evaluate the roles of the *N*- and *C*-termini in receptor binding and to determine the minimal sequence required for effective receptor activation, we conducted sequential truncations of the *Sgg* GSP signal from both ends. We also wanted to identify the residues in KNK-*Sgg* GSP that lead to a ∼100-fold reduction in potency compared to *Sgg* GSP, thus we included in our analysis the sequential truncation of KNK-*Sgg* GSP to *Sgg* GSP. Starting with KNK-*Sgg* GSP, removal of a single residue (K) from the *N*-terminus was sufficient to increase the potency of the peptide by 20-fold, resulting in an analog, NK-*Sgg* GSP, that was only ∼5-fold less potent than *Sgg* GSP (**Table 3**). Removal of an additional residue resulted in an analog, K-*Sgg* GSP, that was as potent as *Sgg* GSP. These results are very intriguing as they indicate that the *N*-terminus of *Sgg* GSP is quite promiscuous, with the exception of the first residue after the double glycine leader, the position most commonly processed in streptococci. Moving to the native *Sgg* GSP, the importance of the *C*-terminus was validated as truncation of even a single residue led to loss of activity (**Table 3**). Contrary to the *C*-terminus and to our surprise, we found that the *N*-terminus region is completely dispensable up to the seventh position (P7) and can accommodate the loss of up to nine residues without a significant reduction in potency (*Sgg* GSP-des-D1-D9 exhibiting an EC_50_ value of 26.9 nM, only 9-fold lower as compared to *Sgg* GSP; **Table 3**). Removal of the tenth residue (W10) resulted in >35-fold reduction in potency compared to the analog lacking nine residues and >330-fold reduction in potency compared to *Sgg* GSP. Interestingly, it appears that removal of entire residues is more tolerated than replacement of the side chains of these residues with that of alanine, suggesting that although the *N*-terminus does not play a significant role in receptor binding, modifications to this region may lead to conformational changes that result in steric clashes and thus significantly lowered potency.

**Table 3.** EC_50_ values of elongated and truncated *Sgg* GSP analogs in UCN34Δ*gsp*^a^

| Compound | Sequence | EC<sub>50</sub> (nM)<sup>b</sup> | 95% CI<sup>c</sup> |
| --- | --- | --- | --- |
| **KNK-*Sgg* GSP** | **KNKDFLIVGPFDWLKKNHKPTKHA** | 287 | 145 – 568 |
| **NK-*Sgg* GSP** | **NKDFLIVGPFDWLKKNHKPTKHA** | 14.1 | 8.01 – 24.7 |
| **K-*Sgg* GSP** | **KDFLIVGPFDWLKKNHKPTKHA** | 2.33 | 1.45 – 3.76 |
| ***Sgg* GSP-des-D1** | **FLIVGPFDWLKKNHKPTKHA** | 9.21 | 4.29 – 19.8 |
| ***Sgg* GSP-des-D1F2** | **LIVGPFDWLKKNHKPTKHA** | 6.49 | 3.03 – 13.9 |
| ***Sgg* GSP-des-D1-L3** | **IVGPFDWLKKNHKPTKHA** | 2.22 | 1.41 – 3.52 |
| ***Sgg* GSP-des-D1-I4** | **VGPFDWLKKNHKPTKHA** | 3.80 | 2.49 – 5.80 |
| ***Sgg* GSP-des-D1-V5** | **GPFDWLKKNHKPTKHA** | 3.60 | 3.34 – 3.89 |
| ***Sgg* GSP-des-D1-G6** | **PFDWLKKNHKPTKHA** | 3.19 | 2.54 – 3.99 |
| ***Sgg* GSP-des-D1-P7** | **FDWLKKNHKPTKHA** | 7.21 | 3.79 – 13.7 |
| ***Sgg* GSP-des-D1-F8** | **DWLKKNHKPTKHA** | 39.7 | 21.4 – 73.6 |
| ***Sgg* GSP-des-D1-D9** | **WLKKNHKPTKHA** | 26.9 | 18.3 – 39.7 |
| ***Sgg* GSP-des-D1-W10** | **LKKNHKPTKHA** | >1000 | -- |
| ***Sgg* GSP-des-A21** | **DFLIVGPFDWLKKNHKPTKH** | >1000 | -- |
| ***Sgg* GSP-des-H20A21** | **DFLIVGPFDWLKKNHKPTK** | >1000 | -- |
| ***Sgg* GSP-des-K19-A21** | **DFLIVGPFDWLKKNHKPT** | >1000 | -- |

<sup>a</sup> See Materials and Methods for details of reporter strains and methods. All assays were performed in triplicate. <sup>b</sup> EC<sub>50</sub> values were determined by testing peptides over a range of concentrations. <sup>c</sup> 95% confidence interval.

## Conclusions

We previously found that *Sgg* produces and secretes a 21-mer peptide that activates bacteriocin production (10). This peptide was named CSP due to its sequence similarity with competence stimulating peptides found in other streptococci. In this work, we showed that the CSP peptide is involved in the activation of gallocin transcription, therefore, we propose to change its name to GSP (gallocin stimulating peptide). GSP displays some unique features since its *N*-terminal amino-acid lies three residues after the double glycine in the leader sequence.

Our results indicate that *Sgg* GSP is secreted by a unique ABC transporter named BlpT and that the SepM-like protease in *Sgg* is not involved in maturation of GSP, but is capable of processing *S. mutans* CSP/MIP to its mature form. Moreover, our SAR studies revealed that Tryptophan at position 10 is important for GSP activity and that nearly half of the GSP signal from the *N*-terminus is entirely dispensable (up to nine residues) for GSP’s ability to activate gallocin expression, while the *C*-terminus is essential for gallocin expression. Our findings provide the groundwork for rationally designing peptide inhibitors targeting gallocin production.

## Acknowledgments

This work was supported in part (Tal-Gan lab) by a grant from the National Science Foundation (CHE-1808370). We want to thank C. R. Bikash (University of Nevada, Reno) for providing purified *S. mutans* 21-CSP and D. G. Cvitkovitch (University of Toronto) for generously providing *S. mutans* SMCOM2 (Δ*comC*, p*comX*::*lacZ*).

## Materials and Methods

All chemical reagents and solvents were purchased from Sigma-Aldrich or Chem-Impex and used without further purification. Water (ddH2O) was purified using a Millipore Analyzer Feed System. Solid-phase Cl-MPA Protide™ resin was purchased from CEM Corporation. 9-Fluorenylmethoxycarbonyl (Fmoc) protected L-α-amino acids were purchased from Advanced ChemTech.

Reverse-phase high-performance liquid chromatography (RP-HPLC) was performed using a Shimadzu UFLC system equipped with a CBM-20A communications bus module, two LC-20AT pumps, a SIL-20A auto sampler, a SPD-20A UV/Vis detector, a CTO-20A column oven, and a FRC-10A fraction collector. All RP-HPLC solvents (ddH_2_O and HPLC-grade acetonitrile (ACN) contained 0.1% trifluoroacetic acid (TFA)). Preparative RP-HPLC was performed using a Phenomenex Kinetex 5 μm 100 Å C18 column (250 x 10 mm) while analytical RP-HPLC was performed using a Phenomenex Kinetex 5 μm 100 Å C18 column (250 x 4.6 mm). Fmoc-based solid-phase peptide synthesis was performed on a Discover Microwave and Liberty1 Automated peptide synthesizer (CEM Corp). Matrix-assisted laser desorption ionization time-of-flight mass spectrometry (MALDI-TOF MS) data were obtained by mixing 0.75 µL of sample with 0.75 µL of matrix solution (α-cyano-4-hydroxycinnamic acid dissolved in ddH_2_O:ACN (1:1) with 0.1% TFA) on a MSP 96 polished steel target plate (Bruker Daltonics) and allowing it to air dry. Data were obtained using a Bruker Microflex spectrometer equipped with a 60 Hz (337 nm wavelength) nitrogen laser and a reflectron. MALDI-TOF MS data were obtained using reflectron positive ion mode with the following settings: ion source 1 = 19 kV, ion source 2 = 15.9 kV, lens = 8.75 kV, reflector = 20 kV, up to 300 Da matrix suppression, 200 laser shots per sample, and detector gain = 1594 V. Exact mass (EM) data were obtained on an Agilent Technologies 6230 time-of-flight mass spectrometer (TOF-MS) with the following settings for positive electrospray ionization mode (ESI+): capillary voltage = 3500 V, fragmentor voltage = 175 V, skimmer voltage = 65 V, octopole RF (Oct 1 RF Vpp) = 750 V, gas temperature = 325 °C, drying gas flow rate = 3 L/min and nebulizer = 25 psi.

### Bacterial growth conditions

The strains used in this study were: *Sgg* UCN34 (wildtype), *Sgg* UCN34 Δ*gallo_rs10340* (Δ*gsp*), *Sgg* UCN34 Δ*gsp* pTCVΩP*blpAB*-*gfp, Sgg* UCN34 Δ*gallo_rs03005* (Δ*sepM*), *Sgg* UCN34 Δ*sepM* pTCVΩP*blpAB*-*gfp, Sgg* UCN34 Δ*gallo_rs10390/10395* (Δ*blpT*), *Sgg* UCN34 Δ*blpT* pTCVΩP*blpAB*-*gfp*. The following procedure was followed for each obtained isolate: a freezer stock was streaked onto a plate of Todd-Hewitt agar supplemented with 0.5% yeast extract (THY plate). Strains containing the pTCVΩP*blpAB*-*gfp* plasmid were grown in the presence of erythromycin (EM) at a final concentration of 10 μg/μL. The plates were incubated for 12-24 h in a CO_2_ incubator (37 °C with 5% CO_2_). Fresh colonies were picked and inoculated into a sterilized culture tube containing 2 mL of sterile THY broth and incubated statically in a CO_2_ incubator overnight. The overnight culture was used for the experiments describe below.

### Solid-phase peptide synthesis

Peptide synthesis and purification of analogs was conducted using previously established methods (10). All peptides were purified to homogeneity (>95%) and their identity validated via mass spectrometry (**Table S1**).

### General assay considerations

Fluorescence and absorbance measurements were recorded using a Biotek Synergy H1 microplate reader using Gen5 data analysis software (v.3.03). Biological assays were performed in triplicate per trial. EC_50_ values from three trials were calculated using GraphPad Prism software (v. 7.0) using a sigmoidal curve fit.

Stock solutions of peptides (1 mM) were prepared in DMSO and stored at 4 °C in sealed glass vials. The maximum concentration of DMSO used in all biological assays did not exceed 2% (v/v). Black polystyrene 96-well microtiter plates (Costar) were used for the GFP cell-based reporter assays.

### Gallocin Induction Reporter Gene Assay Protocol

Peptide stock solutions were serially diluted with DMSO in either 1:2, 1:3 or 1:5 dilutions and 2 μL of the diluted solution was added to each of the wells in a black 96-well microtiter plate. Each concentration was tested in triplicate with DMSO only used as a negative control. Screening of peptide analogs was conducted using the *Sgg* UCN34 Δ*gsp* pTCVΩP*blpAB*-*gfp* strain. An overnight culture was used to make a fresh 2 mL culture without EM using a 1:10 dilution on the day of the experiment and incubated in a CO_2_ incubator (37 °C with 5% CO_2_) for 3-4 hr to reach mid to late log phase of growth (OD_600_ = 0.6 - 0.9). A final 1:10 dilution was made on a larger scale (20 mL) without EM and used for the assay, where 198 μL diluted bacterial culture was added to each well of the microtiter plate containing peptides. Plates were incubated at 37 °C with shaking at 200 rpm. Fluorescence (EX 485 nm and EM 516 nm) and optical density (600 nm) readings were recorded for each well using a plate reader. Measurements were taken 1 hr after inoculation and were recorded every 20-30 min for up to 3 hr to capture the maximum fluorescence signal. The maximum fluorescence signal was normalized with the OD_600_ value and used to construct dose-curves to determine the EC_50_.

### Gallocin Inhibition Screening

Peptide analogs that exhibited less than 50% activation at 10,000 nM relative to 1,000 nM *Sgg* GSP were screened for competitive inhibition. Competitive inhibition was tested by mixing 10,000 nM peptide analog with 20 nM *Sgg* GSP and compared to activation by 20 nM *Sgg* GSP without the addition of synthetic analogs. Peptide analogs were tested in triplicate per trial for three trials (**Data not shown**).

### PDZ domain containing protease/SepM function LC-MS experiment

*Sgg* strains (Δ*gsp*, Δ*sepM*) and *S. mutans* SMCC3 (Δ*comC*) were grown in 60 mL THY overnight at 37 °C with 5% CO_2_ and centrifuged at 4,000 x g for 15 min. The supernatant was discarded, and the cell pellets were washed with 30 mL of sterile saline solution (0.9% w/v NaCl). The cell suspensions were centrifuged at 4,000 x g for 15 min and the supernatant was discarded. The cell pellet was resuspended in 6 mL sterile saline solution and used for the SepM function experiment. The peptides tested: *Sgg* GSP, KNK-*Sgg* GSP and *S. mutans* 21-mer CSP, were resuspended in sterile H_2_O to a final concentration of 1 mM. For each strain tested, around 900 μL of the cell suspension was mixed with 100 μL of each peptide separately to give a 100 μM concentration. “Cell only” control for each strain was made with 900 μL of each cell suspension with 100 μL of sterile saline solution. “Peptide only” control for each peptide was made in sterile saline solution at 100 µM concentration. All variables tested were conducted in sterile 1.5 mL microcentrifuge tubes. All samples were incubated at 37 °C with shaking at 200 rpm for 30 min. After 30 min, the samples were centrifuged at 5,000 x g for 1 min. A 200 μL aliquot from each sample was filter thru a sterile 0.45 μm syringe filter (Phenomenex Phenex™-RC 4mm Syringe Filter) into a sterile 1.5 mL microfuge tube. LC-MS analysis of each sample was performed using XBridge C18 column (5 μm, 4.6 x 150 mm) on an Agilent Technology 1200 series LC connected to Agilent Technologies 6230 TOF-MS. The solvents used for LC-MS were mobile phase A = ddH_2_O + 0.1% formic acid and mobile phase B = ACN + 0.1% formic acid. The LC-MS method used to analyze components in the filtrates had an injection volume of 100 uL and the following linear gradient with a flow rate of 0.5 mL per minute: 5% → 95% B over 25 min.

### LC-MS identifying secreted GSP pheromone

*Sgg* strains (wild-type, Δ*sepM* and Δ*blpT*) were grown in 40 mL of sterile THY at 37 °C with 5% CO_2_ for 12 hrs. The cultures were centrifuged at 4,000 x g for 15 min, the supernatant was discarded, and the cell pellets were resuspended in 5 mL of sterile THY. The resuspended cells were incubated for 16 hrs and centrifuged at 4,000 x g for 15 min. The supernatant was filter-steriled using a 0.22 μm syringe filter (CELLTREAT PES 30 mm diameter Syring Filter) into a sterile 1.5 mL microfuge tube. LC-MS analysis of each sample was performed using XBridge C18 column (5 μm, 4.6 x 150 mm) on an Agilent Technology 1200 series LC connected to Agilent Technologies 6230 TOF-MS. The solvents used for LC-MS were mobile phase A = ddH_2_O + 0.1% formic acid and mobile phase B = ACN + 0.1% formic acid. The LC-MS method used to analyze components in the filtrates had an injection volume of 50 uL and the following linear gradient with a flow rate of 0.5 mL per minute: 5% → 95% B over 25 min.

### Bacteriocin Activity Assay

For well diffusion assays, 2 mL of prey bacteria (here exponentially growing SGM OD_600_ ≈ 0.5) were poured on a THY agar plate. After removal of excess liquid, the plate was dried for 15 min. Using sterile tips, 5-mm-diameter wells were dug into the agar. Wells were filled with 80 μL of *Sgg* filtered supernatant supplemented with 0.1% Tween 20. When the wells were dry, plates were incubated inverted overnight at 37 °C and the inhibition rings were observable the next day.

### Construction of deletion mutants in *Sgg* strain UCN34

The Δ*blpT* (*gallo_rs10390-10395*) and Δ*sepM* (*gallo_rs03005*) mutants were constructed as reported previously in *Sgg* strain UCN34 (33). The primers used are listed in **Table 4**. Briefly, a 1 kb fragment corresponding to the 5’ and 3’ ends of the region to delete was obtained by splicing-by-overlap extension PCR and digested with the restriction enzymes *Eco*RI and *Bam*HI and cloned into the thermosensitive vector pG1-oriT_TnGBS1_. The resulting plasmids pG1Ω*abc* and pG1Ω*sepM* were electroporated in *Streptococcus agalactiae* NEM316 and transferred to *Sgg* UCN34 by conjugation. Chromosomal integration of the plasmid was selected on THY containing erythromycin (10 μg/mL) at 38 °C, a non-permissive temperature for plasmid replication. Then, excision of the plasmid from the chromosome by a second event of homologous recombination was obtained by successive cultures at 30 °C in THY broth without erythromycin resulting in either gene deletion or back to the wild type (bWT) clones. Mutants were systematically tested for erythromycin sensitivity to confirm the loss of the plasmid and confirmed by PCR and sequencing of the chromosomal locus flanking the deletion region.

**Table 4.**
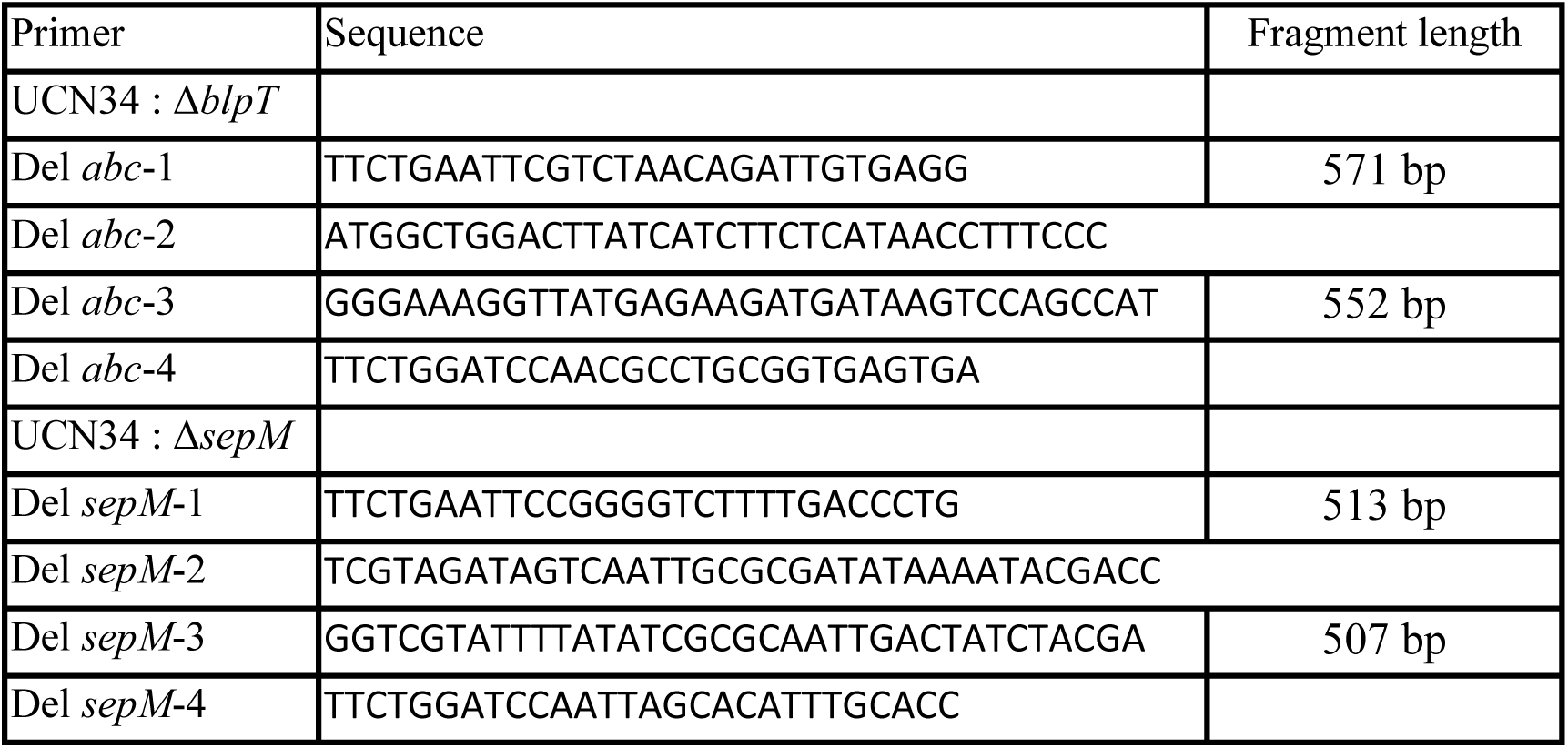
Primers used in this study

### Indirect detection of GSP in the supernatant

Strains tested for GSP production were inoculated at DO_600_=0,1 in THY and grown at 37°C. At DO_600_=1, the cells were pelleted and the supernatants tested for GSP presence were harvested and filtered. After an overnight culture, 100μL of the reporter strain *Sgg* Δ*gsp* pTCVΩP*blpAB*-*gfp* were pelleted and resuspended in 1mL of the tested supernatant. After 3 hours of incubation at 37°C, the fluorescence of the reporter strain was measured by flow cytometry using MACSQuant® VYB flow cytometer.

## Supporting information

### LC-MS identifying secreted GSP pheromone

**Figure S1.**
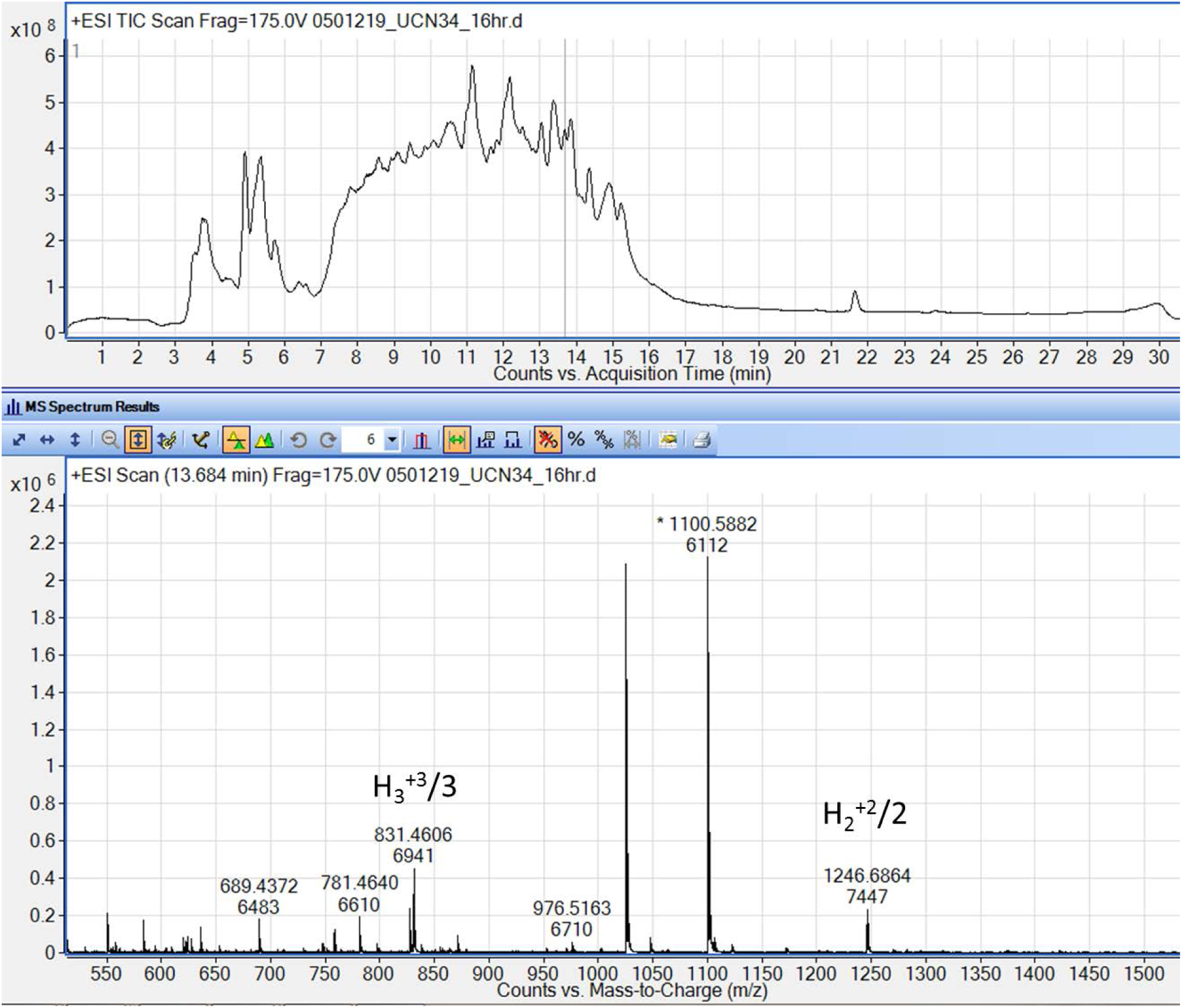
LC-MS of UCN34 supernatant after 16 hours incubation. *Sgg* GSP expected: H_2_^+2^/2 [1246.6855 Da] and H_3_^+3^/3 [831.4594 Da].

**Figure S2.**
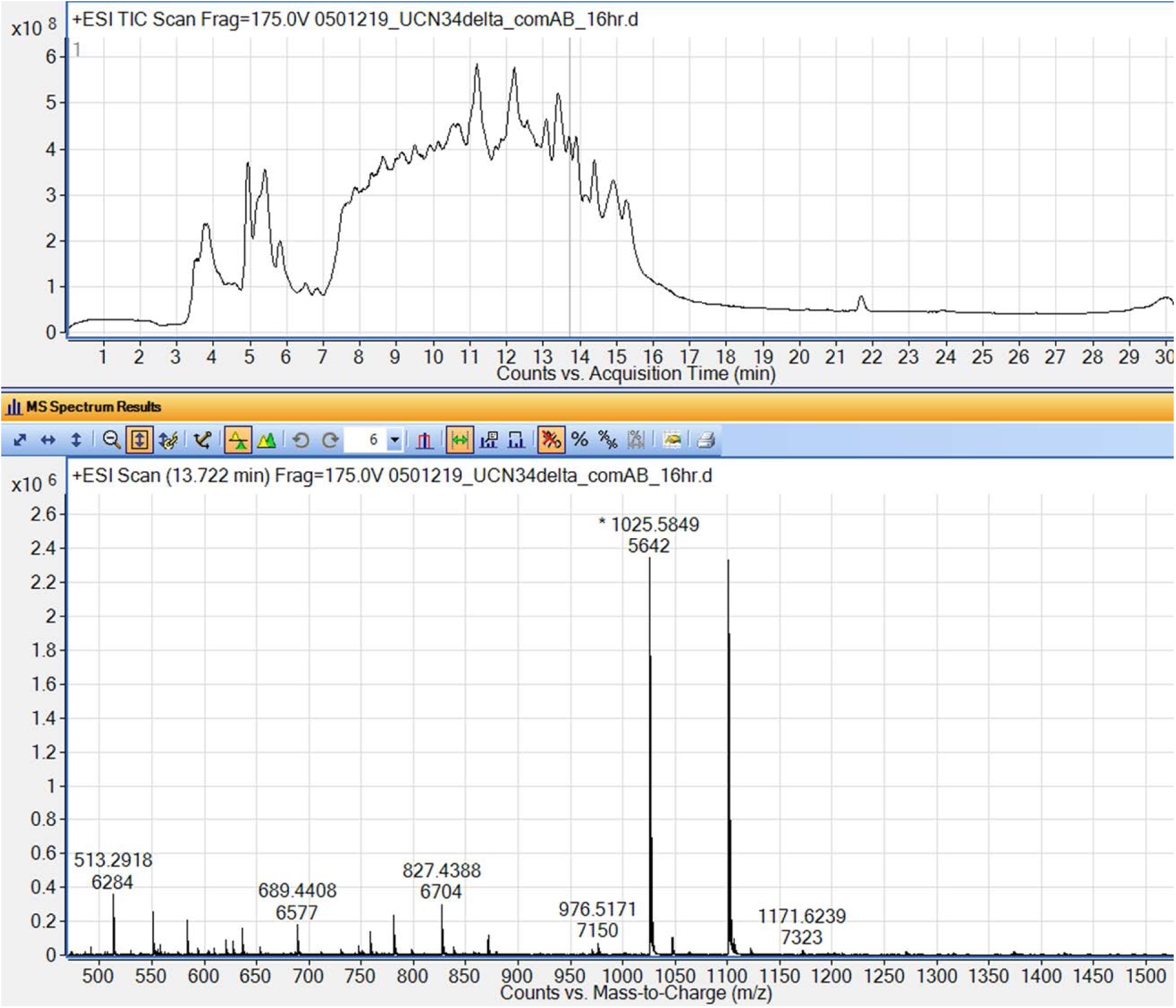
LC-MS of UCN34Δ*blpT* supernatant after 16 hours incubation. No *Sgg* GSP is detected.

**Figure S3.**
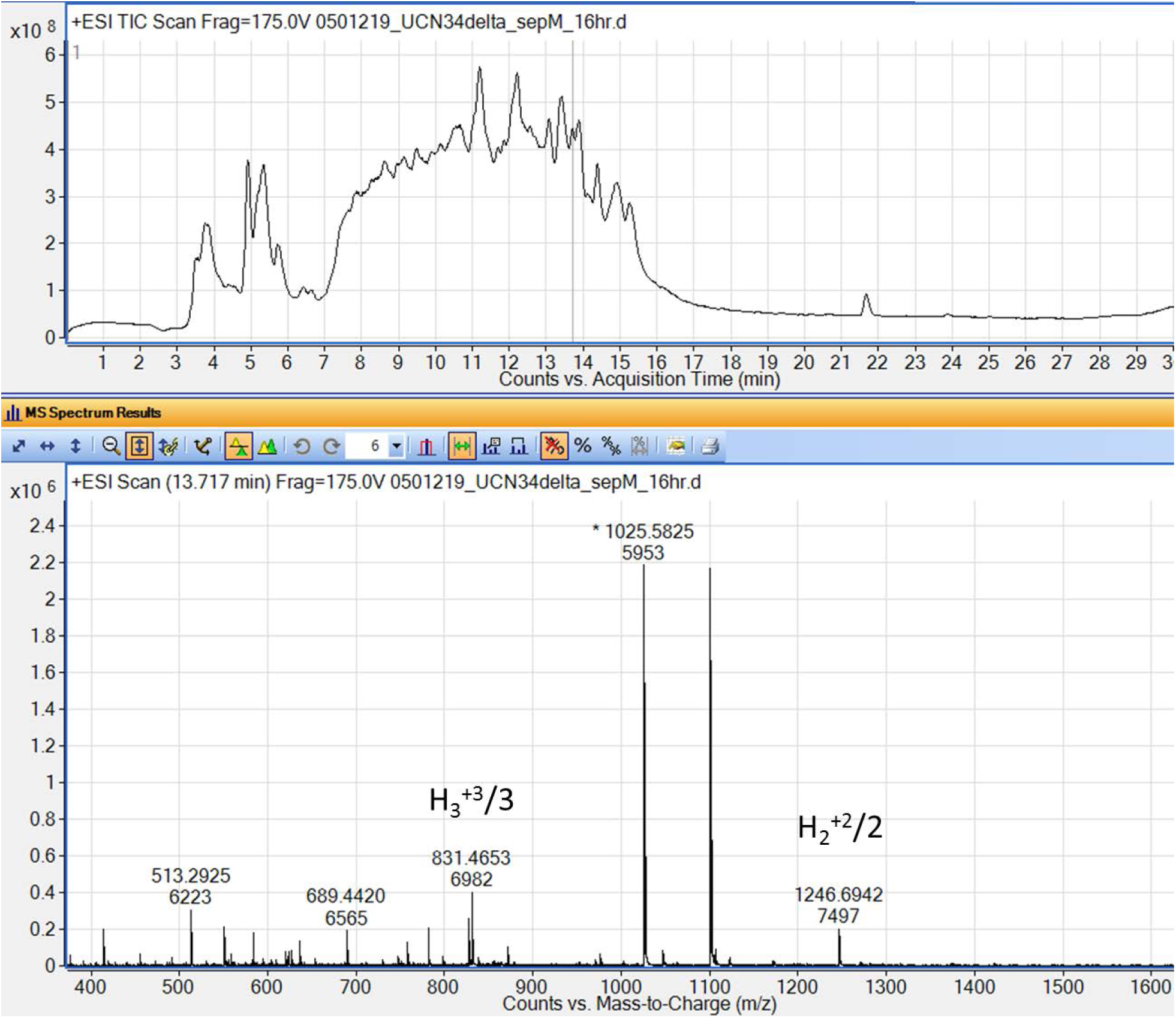
LC-MS of UCN34Δ*sepM* supernatant after 16 hours incubation. *Sgg* GSP expected: H_2_^+2^/2 [1246.6855 Da] and H_3_^+3^/3 [831.4594 Da].

### LC-MS results of SepM Studies

**Figure S4.**
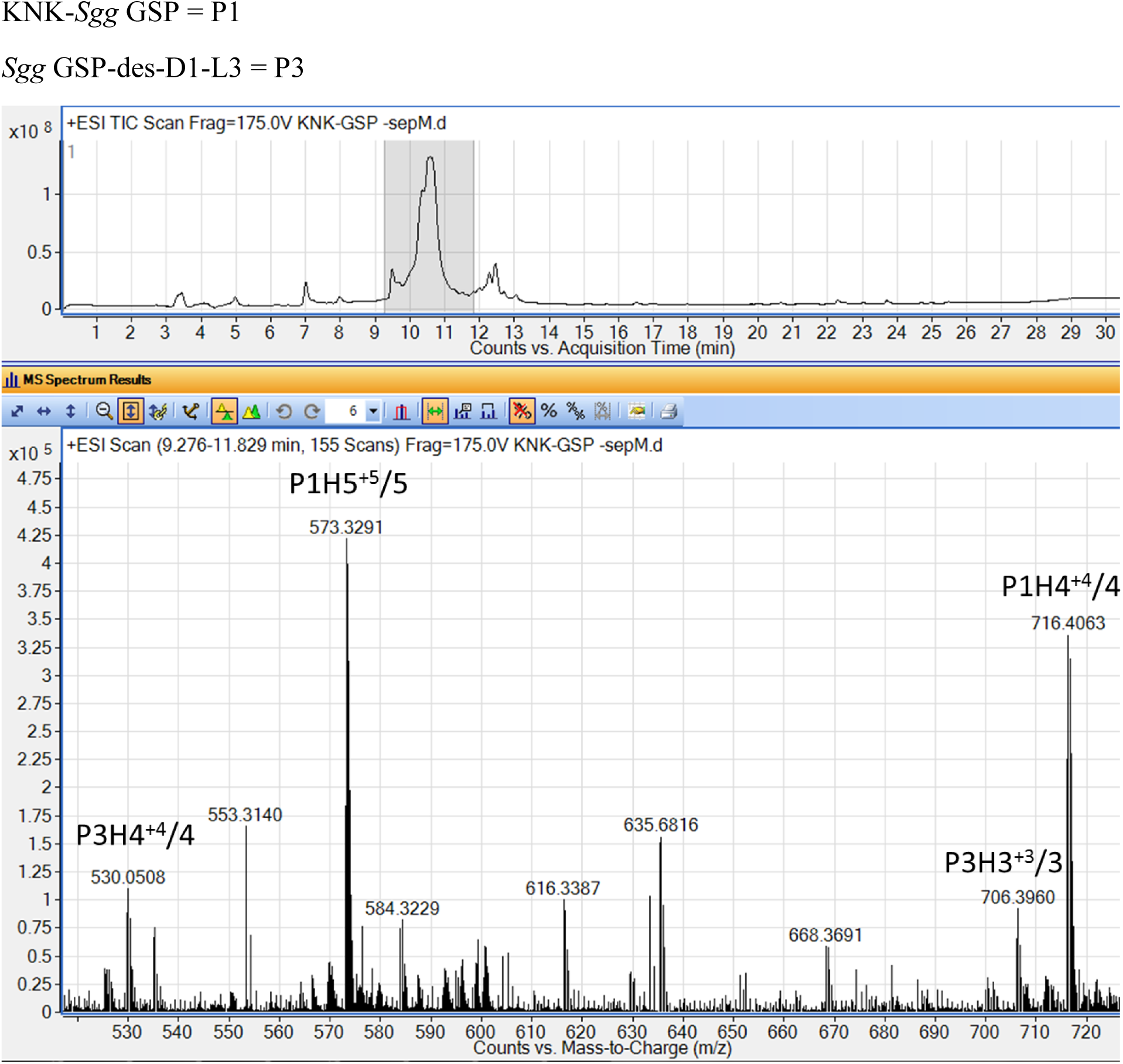
LC-MS of KNK-*Sgg* GSP incubated with UCN34Δ*sepM* cells in saline solution for 30 min. KNK-*Sgg* GSP (P1) expected: P1H_4_^+4^/4 [716.4046 Da] and P1H_4_^+5^/5 [573.3251 Da]. *Sgg* GSP-des-D1-L3 (P3) expected: P3H_3_^+3^/3 [706.3996 Da] and P1H_4_^+4^/4 [530.0515 Da].

**Figure S5.**
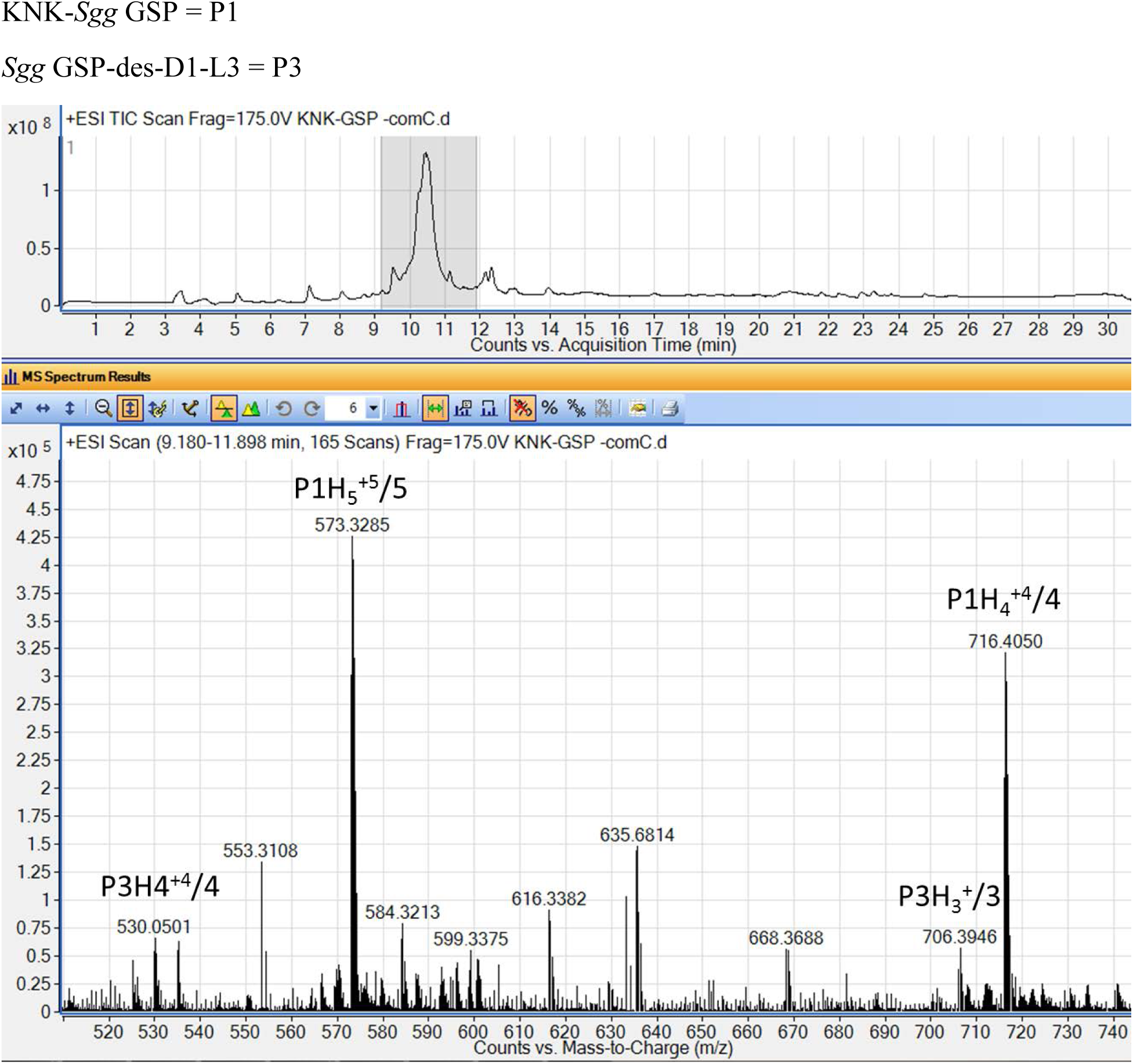
LC-MS of KNK-*Sgg* GSP incubated with UCN34Δ*gsp* cells in saline solution for 30 min. KNK-*Sgg* GSP expected: P1H_4_^+4^/4 [716.4046 Da] and P1H_5_^+5^/5 [573.3251 Da]. *Sgg* GSP-des-D1-L3 expected: P3H_3_^+3^/2 [706.3996 Da] and P3H_4_^+^/4 [530.0515 Da].

**Figure S6.**
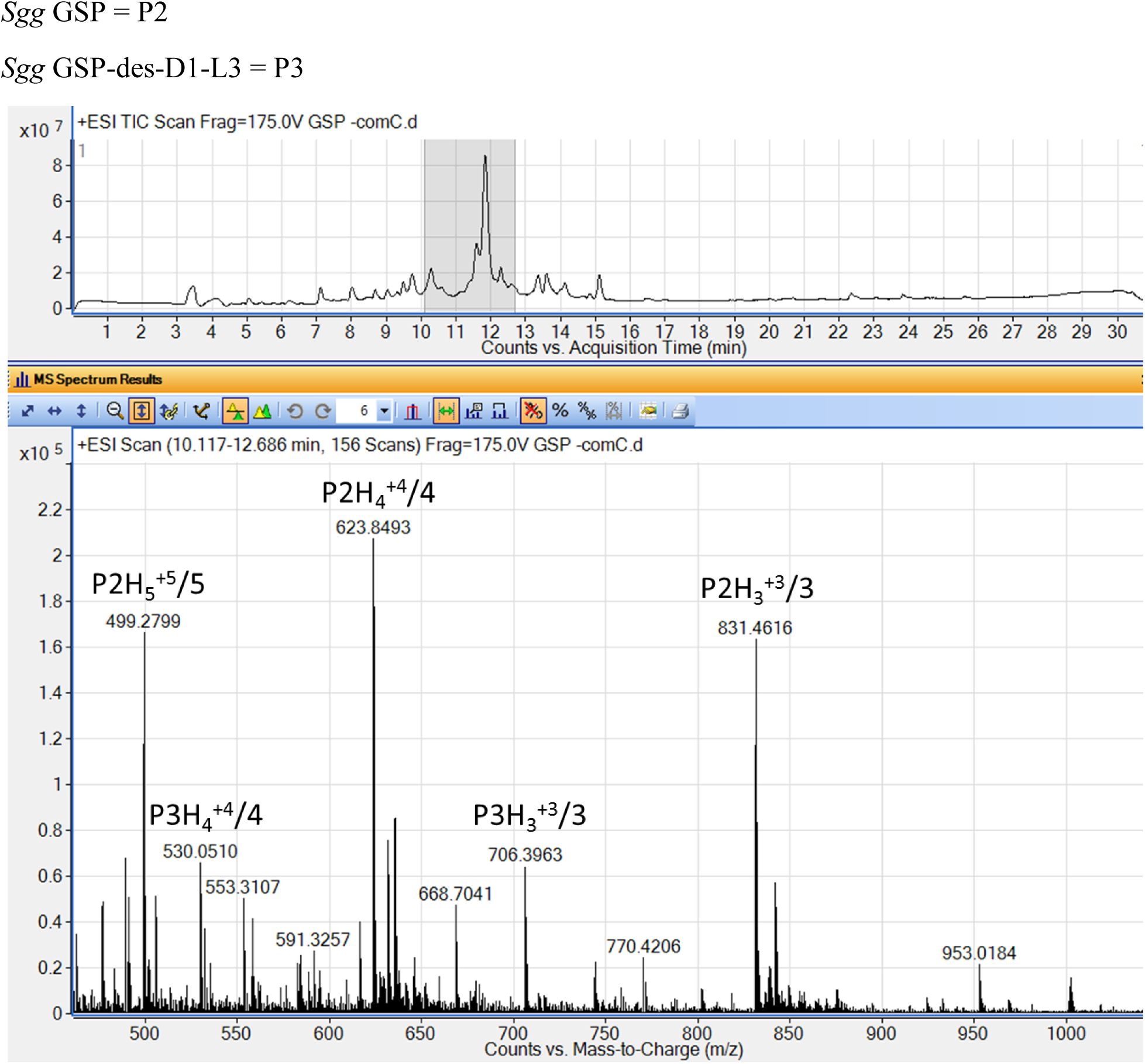
LC-MS of *Sgg* GSP incubated with UCN34Δ*gsp* cells in saline solution for 30 min. *Sgg* GSP (P2) expected: P2H_3_^+3^/3 [831.4594 Da], P2H_4_^+4^/4 [623.8464 Da] and P2H_5_^+^/5 [499.2786 Da]. *Sgg* GSP-des-D1-L3 (P3) expected: P3H_3_^+3^/3 [706.3996 Da] and P3H_4_^+4^/4 [530.0515 Da].

**Figure S7.**
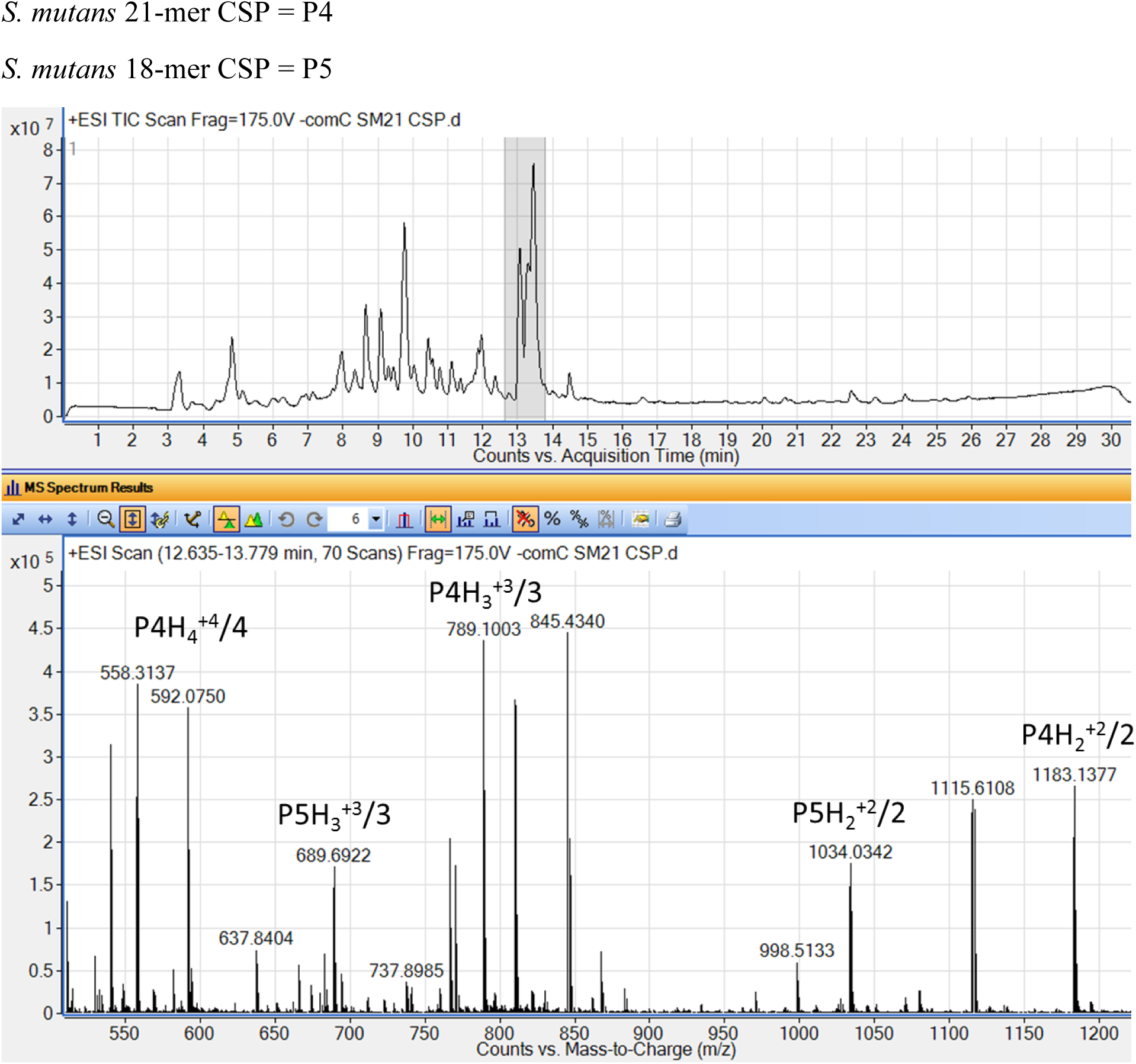
LC-MS of *S. mutans* 21-mer CSP incubated with UCN34Δ*gsp* cells in saline solution for 30 min. *S. mutans* 21-mer (P4) expected: P4H_2_^+2^/2 [1183.128 Da], P4H_3_^+3^/3 [789.0878 Da] and P4H_4_^+4^/4 [592.0676 Da]. *S. mutans* 18-mer CSP (P5) expected: P5H_2_^+2^/2 [1034.0278 Da] and P4H_3_^+3^/3 [689.6876 Da].

**Figure S8.**
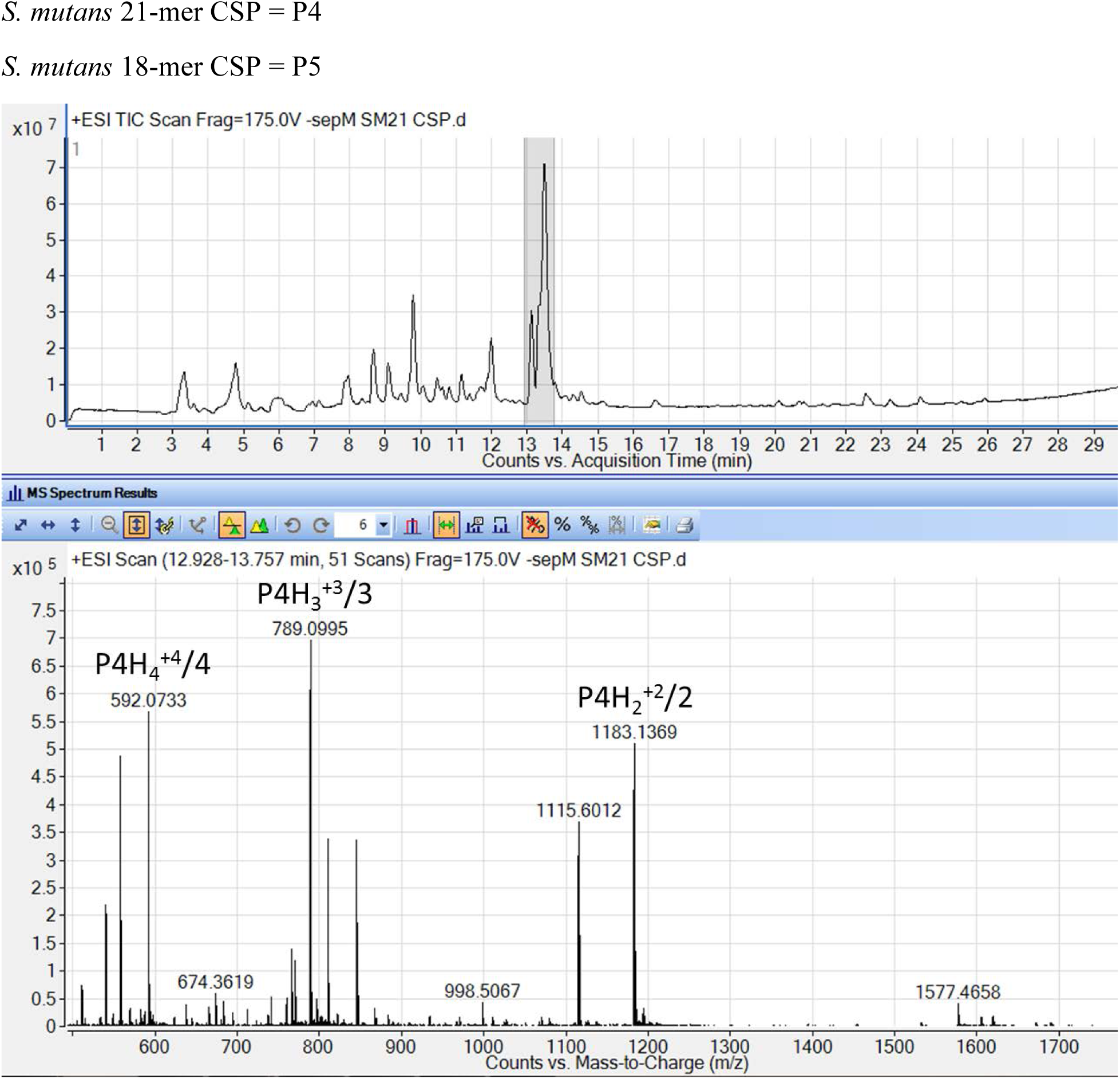
LC-MS of *S. mutans* 21-mer CSP incubated with UCN34Δ*sepM* cells in saline solution for 30 min. *S. mutans* 21-mer (P4) expected: P4H_2_^+2^/2 [1183.128 Da], P4H_3_^+3^/3 [789.0878 Da] and P4H_4_^+4^/4 [592.0676 Da].

**Figure S9.**
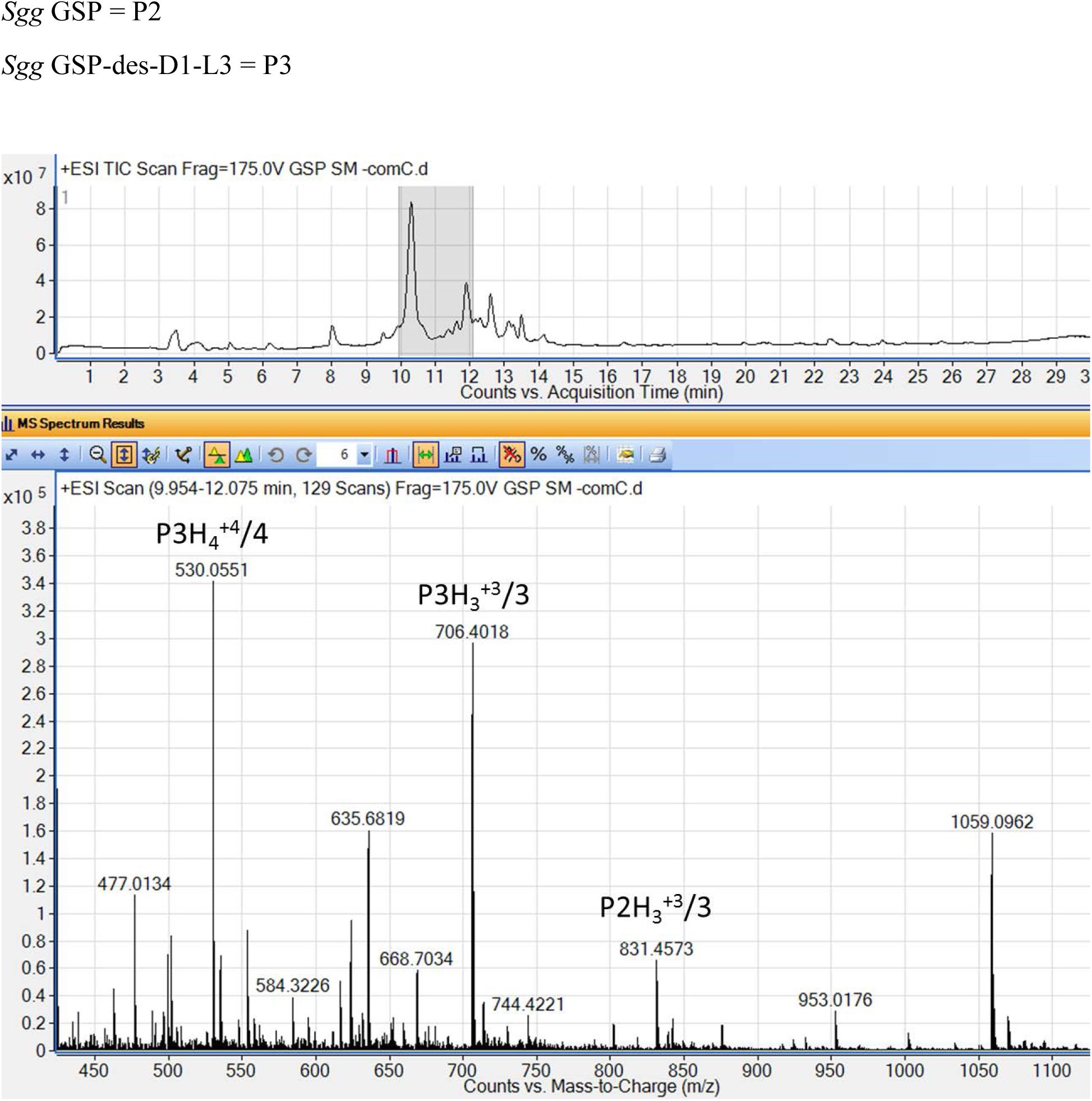
LC-MS of *Sgg* GSP incubated with *S. mutans* Δ*comC* cells in saline solution for 30 min. *Sgg* GSP (P2) expected: P2H_3_^+3^/3 [831.4594 Da]. *Sgg* GSP-des-D1-L3 (P3) expected: P3H_3_^+3^/3 [706.3996 Da] and P3H_4_^+4^/4 [530.0515 Da].

### MS and HPLC data of *Sgg* GSP analogs

**Table S1.** MS and HPLC data of *Sgg* GSP analogs

| Compound Name | Calc. EM [MH<sub>3</sub>]<sup>+3/3</sup> | Obs. EM [MH<sub>3</sub>]<sup>+3/3</sup> | Purity (%) |
| --- | --- | --- | --- |
| *Sgg* GSP D1A | 816.7962 | 816.7961 | > 99 |
| *Sgg* GSP F2A | 806.1157 | 806.1149 | > 96 |
| *Sgg* GSP L3A | 817.4438 | 817.4423 | > 98 |
| *Sgg* GSP I4A | 817.4438 | 817.4420 | > 99 |
| *Sgg* GSP V5A | 822.1157 | 822.1125 | > 99 |
| *Sgg* GSP G6A | 836.1313 | 836.1272 | > 99 |
| *Sgg* GSP P7A | 617.3425 [MH<sub>4</sub>]<sup>+4/4</sup> | 617.3395 [MH<sub>4</sub>]<sup>+4/4</sup> | > 99 |
| *Sgg* GSP F8A | 806.1157 | 806.1132 | > 99 |
| *Sgg* GSP D9A | 816.7962 | 816.7952 | > 99 |
| *Sgg* GSP W10A | 793.1120 | 793.1087 | > 97 |
| *Sgg* GSP L11A | 817.4438 | 817.4406 | > 99 |
| *Sgg* GSP K12A | 812.4402 | 812.4377 | > 98 |
| *Sgg* GSP K13A | 812.4402 | 812.4396 | > 99 |
| *Sgg* GSP N14A | 817.1242 | 817.1244 | > 97 |
| *Sgg* GSP H15A | 809.4522 | 809.4504 | > 99 |
| *Sgg* GSP K16A | 812.4402 | 812.4375 | > 99 |
| *Sgg* GSP P17A | 822.7876 | 822.7867 | > 98 |
| *Sgg* GSP T18A | 616.3438 [MH<sub>4</sub>]<sup>+4/4</sup> | 616.3413 [MH<sub>4</sub>]<sup>+4/4</sup> | > 95 |
| *Sgg* GSP K19A | 609.5819 | 609.5796 | > 99 |
| *Sgg* GSP H20A | 809.4522 | 809.4485 | > 98 |
| KNK-*Sgg* GSP | 716.4046 [MH<sub>4</sub>]<sup>+4/4</sup> | 716.4021 [MH<sub>4</sub>]<sup>+4/4</sup> | > 99 |
| NK-*Sgg* GSP | 912.1721 | 912.1726 | > 98 |
| K-*Sgg* GSP | 874.1578 | 874.1547 | > 97 |
| *Sgg* GSP-des-D1 | 595.0897 [MH<sub>4</sub>]<sup>+4/4</sup> | 595.0869 [MH<sub>4</sub>]<sup>+4/4</sup> | > 97 |
| *Sgg* GSP-des-D1F2 | 558.3226 [MH<sub>4</sub>]<sup>+4/4</sup> | 558.3211 [MH<sub>4</sub>]<sup>+4/4</sup> | > 96 |
| *Sgg* GSP-des-D1-L3 | 706.3996 | 706.3964 | > 99 |
| *Sgg* GSP-des-D1-I4 | 668.7049 | 668.7026 | > 98 |
| *Sgg* GSP-des-D1-V5 | 635.6821 | 635.6801 | > 96 |
| *Sgg* GSP-des-D1-G6 | 616.6750 | 616.6745 | > 99 |
| *Sgg* GSP-des-D1-P7 | 583.9898 | 583.9899 | > 97 |
| *Sgg* GSP-des-D1-F8 | 801.9468 [MH<sub>2</sub>]<sup>+2/2</sup> | 801.9448 [MH<sub>2</sub>]<sup>+2/2</sup> | > 98 |
| *Sgg* GSP-des-D1-D9 | 744.4333 [MH<sub>2</sub>]<sup>+2/2</sup> | 744.4365 [MH<sub>2</sub>]<sup>+2/2</sup> | > 96 |
| *Sgg* GSP-des-D1-W10 | 434.5982 | 434.5997 | > 98 |
| *Sgg* GSP-des-A21 | 807.7804 | 807.7769 | > 96 |
| *Sgg* GSP-des-H20A21 | 762.0941 | 762.0955 | > 98 |
| *Sgg* GSP-des-K19-A21 | 719.3958 | 719.3952 | > 99 |

EM = Exact Mass

## References

1. Schlegel L, Grimont F, Ageron E, Grimont PA, & Bouvet A (2003) Reappraisal of the taxonomy of the Streptococcus bovis/Streptococcus equinus complex and related species: description of Streptococcus gallolyticus subsp. gallolyticus subsp. nov., S. gallolyticus subsp. macedonicus subsp. nov. and S. gallolyticus subsp. pasteurianus subsp. nov. International Journal of Systematic and Evolutionary Microbiology 53(3):631–645.

2. Dekker JP & Lau AF (2016) An update on the Streptococcus bovis group: classification, identification, and disease associations. Journal of clinical microbiology 54(7):1694–1699.

3. Dumke J, et al. (2015) Potential transmission pathways of Streptococcus gallolyticus subsp. gallolyticus. PLoS One 10(5):e0126507.

4. Boleij A, van Gelder MM, Swinkels DW, & Tjalsma H (2011) Clinical Importance of Streptococcus gallolyticus infection among colorectal cancer patients: systematic review and meta-analysis. Clinical Infectious Diseases 53(9):870–878.

5. Boleij A, et al. (2011) Novel clues on the specific association of Streptococcus gallolyticus subsp gallolyticus with colorectal cancer. Journal of Infectious Diseases 203(8):1101–1109.

6. Tjalsma H, Boleij A, Marchesi JR, & Dutilh BE (2012) A bacterial driver–passenger model for colorectal cancer: beyond the usual suspects. Nature Reviews Microbiology 10(8):575.

7. Geng J, et al. (2014) Co-occurrence of driver and passenger bacteria in human colorectal cancer. Gut pathogens 6(1):26.

8. Kumar R, et al. (2017) Streptococcus gallolyticus subsp. gallolyticus promotes colorectal tumor development. PLoS pathogens 13(7):e1006440.

9. Aymeric L, et al. (2018) Colorectal cancer specific conditions promote Streptococcus gallolyticus gut colonization. Proceedings of the National Academy of Sciences of the United States of America 115(2):E283–E291.

10. Harrington A & Tal-Gan Y (2018) Identification of Streptococcus gallolyticus subsp. gallolyticus (biotype I) competence stimulating peptide pheromone. Journal of bacteriology:JB. 00709-00717.

11. Tomasz A (1966) Model for the Mechanism Controlling the Expression of Competent State in Pneumococcus Cultures. Journal of Bacteriology 91(3):1050–1061.

12. Håvarstein LS, Coomaraswamy G, & Morrison DA (1995) An unmodified heptadecapeptide pheromone induces competence for genetic transformation in Streptococcus pneumoniae. Proceedings of the National Academy of Sciences of the United States of America 92(24):11140–11144.

13. Havarstein LS, Diep DB, & Nes IF (1995) A family of bacteriocin ABC transporters carry out proteolytic processing of their substrates concomitant with export. Molecular microbiology 16(2):229–240.

14. Shanker E & Federle MJ (2017) Quorum Sensing Regulation of Competence and Bacteriocins in Streptococcus pneumoniae and mutans. Genes 8(1):15.

15. Luo P, Li H, & Morrison DA (2003) ComX is a unique link between multiple quorum sensing outputs and competence in Streptococcus pneumoniae. Molecular microbiology 50(2):623–633.

16. Håvarstein LS, Hakenbeck R, & Gaustad P (1997) Natural competence in the genus Streptococcus: evidence that streptococci can change pherotype by interspecies recombinational exchanges. Journal of bacteriology 179(21):6589–6594.

17. Ween O, Teigen S, Gaustad P, Kilian M, & Håvarstein LS (2002) Competence without a competence pheromone in a natural isolate of Streptococcus infantis. Journal of bacteriology 184(13):3426–3432.

18. Tong H, et al. (2006) Establishing a genetic system for ecological studies of Streptococcus oligofermentans. FEMS microbiology letters 264(2):213–219.

19. Fontaine L, et al. (2010) A novel pheromone quorum-sensing system controls the development of natural competence in Streptococcus thermophilus and Streptococcus salivarius. Journal of bacteriology 192(5):1444–1454.

20. Mashburn-Warren L, Morrison DA, & Federle MJ (2010) A novel double-tryptophan peptide pheromone controls competence in Streptococcus spp. via an Rgg regulator. Molecular microbiology 78(3):589–606.

21. Wang CY & Dawid S (2018) Mobilization of Bacteriocins during competence in Streptococci. Trends in microbiology 26(5):389–391.

22. Ahn S-J, Wen ZT, & Burne RA (2006) Multilevel Control of Competence Development and Stress Tolerance in <em>Streptococcus mutans</em> UA159. Infection and Immunity 74(3):1631–1642.

23. Li Y-H, Lau PC, Lee JH, Ellen RP, & Cvitkovitch DG (2001) Natural genetic transformation ofstreptococcus mutans growing in biofilms. Journal of bacteriology 183(3):897–908.

24. Li Y-H, et al. (2002) A quorum-sensing signaling system essential for genetic competence in Streptococcus mutans is involved in biofilm formation. Journal of bacteriology 184(10):2699–2708.

25. Lemme A, Gröbe L, Reck M, Tomasch J, & Wagner-Döbler I (2011) Subpopulation-specific transcriptome analysis of competence-stimulating-peptide-induced Streptococcus mutans. Journal of bacteriology 193(8):1863–1877.

26. Reck M, Tomasch J, & Wagner-Döbler I (2015) The alternative sigma factor SigX controls bacteriocin synthesis and competence, the two quorum sensing regulated traits in Streptococcus mutans. PLoS genetics 11(7):e1005353.

27. Premnath P, Reck M, Wittstein K, Stadler M, & Wagner-Döbler I (2018) Screening for inhibitors of mutacin synthesis in Streptococcus mutans using fluorescent reporter strains. BMC microbiology 18(1):24.

28. Biswas S, Cao L, Kim A, & Biswas I (2016) SepM, a streptococcal protease involved in quorum sensing, displays strict substrate specificity. Journal of bacteriology 198(3):436–447.

29. Hossain MS & Biswas I (2012) An extracelluar protease, SepM, generates functional CSP in Streptococcus mutans UA159. Journal of bacteriology:JB. 01381–01312.

30. Fontaine L, et al. (2013) Mechanism of competence activation by the ComRS signalling system in streptococci. Molecular microbiology 87(6):1113–1132.

31. Lin I-H, et al. (2011) Sequencing and comparative genome analysis of two pathogenic Streptococcus gallolyticus subspecies: genome plasticity, adaptation and virulence. PLoS One 6(5):e20519.

32. Morrison DA, Guédon E, & Renault P (2013) Competence for natural genetic transformation in the Streptococcus bovis group streptococci S. infantarius and S. macedonicus. Journal of bacteriology 195(11):2612–2620.

33. Danne C, Guerillot R, Glaser P, Trieu-Cuot P, & Dramsi S (2013) Construction of isogenic mutants in Streptococcus gallolyticus based on the development of new mobilizable vectors. Res Microbiol 164(10):973–978.

